# Inducible volatile chemical signalling drives antifungal activity of *Trichoderma hamatum* GD12 during confrontation with the pathogen *Sclerotinia sclerotiorum*

**DOI:** 10.1101/2025.07.07.662534

**Authors:** Gareth A. Thomas, József Vuts, David M. Withall, John C. Caulfield, John Sidda, Murray R. Grant, Christopher R. Thornton, Michael A. Birkett

**Affiliations:** Protecting Crops and the Environment, Rothamsted Research, Harpenden, AL5 2JQ; Biosciences, College of Life and Environmental Sciences, University of Exeter, Exeter, EX4 4QD; School of Life Sciences, University of Warwick, Coventry, CV4 7AL

**Keywords:** *Trichoderma*, *Sclerotinia sclerotiorum*, volatile organic compounds, antagonism, 1-octen-3-one

## Abstract

**BACKGROUND:** The use of beneficial soil fungi or their natural products offers a more sustainable alternative to synthetic fungicides for pathogen management in crops. Volatile organic compounds (VOCs) produced by such fungi act as semiochemicals that inhibit pathogens, with VOC production influenced by physical interactions between competing fungi. This study explores the interaction between the beneficial soil fungus *Trichoderma hamatum* GD12 strain (GD12), previously shown to antagonize crop pathogens such as *Sclerotinia sclerotiorum*, to test the hypothesis that its antagonistic effect is mediated by volatile chemical signalling. A GD12 mutant deficient in the chitinolytic enzyme *N*-acetyl-β-glucosaminidase (*ΔThnag : : hph*), which shows reduced biocontrol activity, was also examined.

*RESULTS:* In dual-culture confrontation assays, co-inoculation of GD12 and *S. sclerotiorum* led to fungistatic interactions after 7 days, whereas *ΔThnag : : hph* showed no antagonism, indicating a loss of antagonistic function. VOCs collected from individual and co-cultures were analysed by gas chromatography – flame ionization detector (GC-FID) analysis and coupled GC-mass spectrometry (GC-MS), revealing significant differences in VOC production between treatments, with VOC production notably upregulated in the GD12 + *S. sclerotiorum* co-culture. Peak production of 6-pentyl-2H-pyran-2-one occurred 17 days post-inoculation. This upregulation was absent in the *ΔThnag : : hph* co-culture, suggesting VOCs may drive antagonism. Synthetic VOC assays revealed several compounds inhibitory to *S. sclerotiorum*, including 1-octen-3-one, which also arrested the growth of key fungal pathogens (*Botrytis cinerea*, *Pyrenopeziza brassicae*, and *Gaeumannomyces tritici*). Structural insights into 1-octen-3-one’s antifungal activity against *S. sclerotiorum* are also presented.

*CONCLUSIONS:* These findings support the hypothesis that the antagonistic properties of *T. hamatum* GD12 against crop fungal pathogens can, in part, be attributed to VOC production. Further research is needed to assess the potential of these semiochemicals as tools for pathogen management in agriculture.

## Introduction

*Sclerotinia sclerotiorum* (Lib.) de Bary (Family: Sclerotiniaceae) is a ubiquitous soil-borne fungal pathogen, affecting approximately 800 plant species worldwide, including economically important agricultural crops such as carrots, lettuce, sunflower, oilseed rape and potato (Boland *et al*., 1994; Bolton *et al*., 2006a). Management of *S. sclerotiorum* on agricultural crops relies mainly on the application of synthetic fungicides (Derbyshire and Denton-Giles, 2016), although the over-application of fungicides has increased selective pressure, leading to an increase in frequency of fungicide-resistant strains (Ma *et al*., 2009). Alternative methods for controlling *S. sclerotiorum* include the use of crop rotations, which may also be ineffective due to the formation of vegetative sclerotia by *S. sclerotiorum,* which can remain viable in the soil for over eight years and are resistant to physical, chemical and biological degradation (Tribe, 1957; Adams, 1979; Bolton *et al*., 2006a). Moreover, engineering crop resistance towards the pathogen has also proven challenging due to differing pathovars of the pathogen, and a lack of resistance in major crops, making breeding programmes a challenge (Bolton et al., 2006; Derbyshire et al., 2022). Therefore, more sustainable approaches for pathogen management on crops, that minimise reliance on synthetic fungicides, are needed.

Sustainable management strategies for the control of *S. sclerotiorum* include the exploitation of microbial biocontrol agents of the pathogen, including beneficial soil fungi. These decrease the negative potential of pathogens on crops through direct antagonism of the pathogen, competition for resources (e.g. nutrients), or through modification of plant defence responses (Ghorbanpour *et al*., 2018). *Trichoderma* (Hypocreaceae) is a well-studied genus of beneficial soil fungi due to their ability to inhibit fungal pathogen development, induce plant defence responses against pathogens and promote plant growth (Druzhinina *et al*., 2011; Woo *et al*., 2023). GD12, a strain of *T. hamatum*, is effective at suppressing the growth of *S. sclerotiorum* in peat-based microcosms (Ryder *et al*., 2012 Studholme *et al*., 2013; Shaw *et al*., 2016), with the suppressing capability of GD12 requiring the chitinase gene *N*-acetyl-β-glucosaminidase (Ryder et al., 2012). Genome sequencing of *Trichoderma hamatum* (Feng *et al*., 2025) and specifically GD12 (Studholme et al., 2013) reveals the presence of silent gene clusters which could be activated in the presence of antagonistic microorganisms in soil, leading to the production of secondary metabolites which are not produced under standard laboratory conditions (Shaw et al., 2016). This antagonism can stimulate the induction of secondary metabolite biosynthetic gene clusters as evidenced by induction of genes encoding predicted polyketide synthases (PKSs) and Non-Ribosomal Peptide Synthetases (NRPSs) clusters during the interaction between *S. sclerotiorum* and *T. hamatum* in peat microcosms (Shaw *et al*., 2016). However, the causal metabolites involved in these interactions are currently unknown.

Volatile organic compounds (VOCs) are a class of low molecular weight secondary metabolites produced by soil microorganisms, which contribute to their ability to compete against neighbouring organisms for resources in soil (Garbeva and Weisskopf, 2020; Weisskopf *et al*., 2021). The ability of VOCs to travel between gas- and water-filled pockets in soil classifies them as long-distance messengers, compared to non-volatile secondary metabolites, which may be drivers of more local interactions (Kai *et al*., 2016; Schulz-Bohm *et al*., 2017; Westhoff *et al*., 2017). Microbial VOCs are involved in a range of biological activities, including direct inhibition of pathogenic microorganisms (Fernando *et al*., 2005), induced plant defence against pathogens (Ryu *et al*., 2004) and plant growth promotion (Ryu *et al*., 2003). Several studies indicate *Trichoderma* VOCs can specifically play an inhibitory role against a range of fungal pathogens (Amin *et al*., 2010; Stoppacher *et al*., 2010; Jeleń *et al*., 2014a; Meena *et al*., 2017; Wonglom *et al*., 2020). These biological activities highlight the potential for microbial VOCs to be used as effective alternatives to pesticides and fertilisers (Thomas *et al*., 2020, 2023).

Whilst beneficial soil microbes can produce VOCs when grown axenically under standard laboratory conditions, they exist in complex communities within the soil matrix. Genome sequencing of fungal species indicates that many secondary metabolite gene clusters are silent and telomeric under standard laboratory conditions and may require specific cultivation conditions to activate them, including stress inducing, or co-culturing with different species of microorganisms (Scherlach and Hertweck, 2021). Experimentally reproducing a more natural microbe interaction environment is feasible through inoculating different species of microorganisms within the same confined space. Such an approach may stimulate the antagonism that activate silents gene clusters, and hence facilitate the discovery of novel, bioactive compounds. It has been observed that ornesillic acid production was uniquely induced through co-culturing of *Streptomyces* and *Aspergillus* (Schroeckh *et al*., 2009). This study subsequently led to an expansion in the discovery of novel secondary metabolite production through microbial co-culture (Knowles *et al*., 2022). For example, in *T. harzianum,* co-culture with the endophyte *Talaromyces pinophilus* led to changes in secondary metabolite production relative to monoculture controls (Vinale *et al*., 2017). The majority of these studies focus on changes in non-volatile compound production during physical interactions, however a growing body of evidence suggests co-culturing can also induce VOC production, with examples from fungal-fungal (Hynes *et al*., 2007a; Evans *et al*., 2008; El Ariebi *et al*., 2016; Guo *et al*., 2019a; O’Leary *et al*., 2019), bacterial-bacterial (Tyc *et al*., 2015, 2017) or fungal-bacterial (Albarracín Orio *et al*., 2020) interactions.

Here, we aimed to determine the role of VOCs in the biocontrol capabilities of *T. hamatum* GD12 against *S. sclerotiorum*. We demonstrate; i) quantitative and qualitative changes in VOC production by *T. hamatum* occur during confrontation with *S. sclerotiorum*; ii) temporal changes in VOC production during confrontation occur, with maximal induction day 17 post inoculation; iii) VOCs produced by *T. hamatum* have antifungal activity against *S. sclerotiorum*; iv) identification of 1-octen-3-one, which completely inhibits the growth of *S. sclerotiorum* as well as other agriculturally important fungal pathogens; and iv) the structural features required for the antifungal activity of 1-octen-3-one against *S. sclerotiorum*. This work highlights the power of using *Trichoderma*-pathogen co-culture to reveal cryptic chemistries encoding bioactive VOCs for use as pathogen management tools in agriculture.

## Materials and methods

### Dual-culture confrontation assays. Trichoderma hamatum

GD12 isolated from a potato field (Great Down, Devon, UK), (Thornton, pers. comm.), *ΔThnag : : hph* : : *hph* (Ryder et al., 2012) and *Sclerotinia sclerotiorum* isolate 1 (isolated on oilseed rape petal, ADAS Rosemaund, Herefordshire, UK), (West, pers. comm.), used in the study were maintained on Potato Dextrose Agar (PDA) (15 g Bacteriological Agar No. 2, LabM, UK; 24 g Potato Dextrose Broth (PDB), Sigma, UK; 1000 mL distilled H2O) slopes in sterile, screw-capped plastic vials (ThermoScientific, UK). Circular plugs (5 mm diam.) were cut using a sterilised cork-borer (Sigma, UK) from the leading edge of mycelia of 3-day old PDA plates of *T. hamatum* (GD12, *ΔThnag : : hph*) or *S. sclerotiorum* isolate 1. Each experiment comprised (a) a control, containing uninoculated growth media (PDA), (b) the confrontation of a *T. hamatum* strain co-cultured against itself (self-challenged), (c) the *S. sclerotiorum* co-cultured against itself (self-challenged), and (d) the *T. hamatum* strain challenged against *S. sclerotiorum* (co-culture) (n=4). Plugs from individual strains of *T. hamatum* were placed approximately 80 mm away from plugs of *S. sclerotiorum* on fresh PDA plates (90 mm) and grown for 7 days under a 16 h/8 h fluorescent light/dark photoperiod at 24°C until required for dynamic headspace collection experiments (detailed below).

### Dynamic headspace collection (air entrainment)

PDA plates containing 7-day-old fungal cultures (see above) were enclosed individually in glass entrainment vessels (12 cm diam. x 6 cm height). Charcoal-purified air (flowrate 600 mL/min) was pushed into each entrainment vessel and drawn (flowrate 500 mL/min), ensuring a positive pressure (100 mL/min) throughout the system. Air was drawn through a glass tube containing Porapak Q (50 mg, 50/80 mesh, Supelco, Bellefonte, PA) held with two plugs of silanized glass wool, for 20 h at ambient temperature (Pye volatile collection kits, Kings Walden, UK). Before each collection, glass vessels were washed with Teepol detergent, rinsed with distilled water, washed with acetone (ThermoFisher, UK), and then placed in a modified heating oven (180°C) for a minimum of 2 h. Charcoal filters (10-14 mesh, 50 g) (Sigma, UK) were conditioned prior to each experiment by attaching them to a supply of nitrogen in a modified heating oven (150°C) under a constant stream of nitrogen. Volatile collections were performed under a 16 h/8 h light/dark photoperiod at 24°C. Porapak Q traps were cleaned by washing with freshly redistilled diethyl ether (2 mL) and heated to 132°C for a minimum of 2 h under a stream of nitrogen. Following collections, VOCs were eluted from the Porapak Q traps with freshly redistilled diethyl ether (750 µL) into 1.1 mL pointed vials (ThermoScientific, Germany), capped with an 8 mm Chromacol screw cap vial lid (ThermoScientific, Germany) with an 8 mm Silicone Red PTFE Septa (Kinesis, UK). The eluent was concentrated to 50 µL under a gentle stream of nitrogen and stored at -20°C prior to further analysis.

### Time-course VOC collection experiment

Solid-phase microextraction (SPME) was selected as the method for VOC analysis for time-course experiments rather than dynamic headspace collection, as preliminary experiments showed repeated dynamic headspace collections of VOCs from fungal cultures led to drying out of growth media, impacting fungal growth. An SPME (100 µM Polydimethylsiloxane (PDMS) fibre, Supelco, UK) was introduced into the GC thermal desorption injector port to desorb for 10 min (temperature of injector=250°C). The SPME fibre was inserted through a clean septum and exposed to the headspace of the fungal culture within a clean glass entrainment vessel (12 cm diam. x 6 cm height) for 1 h. SPME samples were taken at 1, 2, 3, 4, 5, 6,7, 10, 17 and 24 days post inoculation (dpi) from cultures of (a) self-challenged *S. sclerotiorum* isolate 1, 1. (b) self-challenged *T. hamatum* GD12, or (c) GD12 co-cultured with *S. sclerotiorum*. The first sampling timepoint (day 1 post inoculation) of the experiment was used as a baseline, with 80 mm of distance between the fungal mycelia. At day 2, the distance between the self-challenged GD12 treatments was 19-24 mm, and 7-15 mm for the GD12 co-cultured with *S. sclerotiorum* treatments. Self-challenged *S. sclerotiorum* treatments had already initiated contact by this stage of sampling. By day 3, contact between mycelia across all treatments had established.

### Gas chromatography – flame ionization detector (GC-FID) analysis

Air entrainment samples were analysed on an Agilent 6890 GC equipped with a cool on-column injector, an FID and a HP-1 bonded-phase fused silica capillary column (50 m x 0.32 mm i. d. x 0.52 µm film thickness). The oven temperature was set at 30°C for 0.1 min, then increased at 5°C/min to 150°C for 0.1 min, then at 10°C/min to 230°C for a further 25 min. The carrier gas was hydrogen. VOCs adsorbed on SPME fibres were thermally desorbed by inserting the fibre directly into the OPTIC Programmable Temperature Vaporisor (PTV) unit (30 -> 250*°*C ballistically at a rate of 16*°*C/s).

### Coupled GC-mass spectrometry (GC-MS)

An Agilent Mass Selective Detector (MSD) 5973 coupled to an Agilent 6890N GC (fitted with a non-polar HP1 column 50 m length x 0.32 mm i. d. x 0.52 µM film thickness, J & W Scientific) was used for analysis. Sample injection was via cool-on column and MS ionization was by electron impact at 70 eV at 220°C. The GC oven temperature was maintained at 30°C for 5 min and then programmed at 5°C/min to 250°C, run time 70 minutes. Tentative identifications were confirmed by co-injections with authentic standards (Pickett et al., 1990).

### Synthetic compound assays

Synthetic standards of VOCs identified from air entrainments of *T. hamatum* GD12 were applied to sterile qualitative filter paper (6 mm) (Whatman, UK) and placed onto a Petri dish containing PDA. On a fresh plate of PDA, a plug (5 mm diam.) of *S. sclerotiorum* was inoculated and the plate inverted over another plate containing the filter paper with the synthetic VOC sample, and the two plates were then sealed with tape, ensuring no physical contact between the VOC sample and the pathogen (Figure S1). Solutions of VOCs were prepared in freshly redistilled diethyl ether to ensure that 20 µL application of solution gave a dose of 45.5 µM. This dose was decided based on preliminary experiments, where 5 µL of each compound were applied neat to a sterile filter paper, and the dose selected based on the least inhibitory compound (1-pentanol; 5 µL of which equates to 45.5 µM). This dose is similar to that used in previous studies (Tyc *et al*., 2015). For compounds demonstrating significant (p < 0.05) antifungal activity relative to control treatments when applied at 45.5 µM, doses onto filter discs were diluted to 22.75 µM, 11.125 µM, 4.55 µM, 2.275 µM, 0.91 µM and 0.455 µM in freshly redistilled diethyl ether from a stock solution, until no further inhibition was observed. Mycelial measurements from *S. sclerotiorum* were taken using a 30 cm ruler after 72 h. From the diameter of *S. sclerotiorum*, the area of colony was calculated as πr^2^, where “r” is equal to the radius of the colony. Control plates contained a sterile filter paper disc with 20 µL of freshly redistilled diethyl ether alone.

### Statistical analysis

For comparison of GC analysis from confrontation assays, peak area values were individually measured (Agilent Chemstation) and log10-transformed. An adjustment of 0.001 was applied to account for values recorded as zero. Statistical comparison of compounds present in both mono- and co-culture treatments were analysed using an unpaired Student’s t-test assuming equal variances, one variate with grouped factor. To establish the antifungal activity of selected VOCs, mycelial areas were statistically compared across treatments using one way analysis of variance (ANOVA), followed by Tukey’s Honest Significant Difference test at the *p* < 0.05 level, where multiple comparisons were required. Genstat^®^ (v21, ©VSN International, Hemel Hempstead, UK) was used for all statistical analyses.

## Results

Dual-culture confrontation assays demonstrated fungistatic interactions between *T. hamatum* GD12 when confronted with *S. sclerotiorum* after 7 days of growth (Figure 1). This was accompanied by the formation of yellow spores by *T. hamatum* GD12 in the interaction zone between the two fungi. Contrastingly, during confrontation with *T. hamatum ΔThnag : : hph*, overgrowth of *S. sclerotiorum* was observed, depicted by a black arrow (Figure 1f), indicating a loss in the antagonistic capabilities of the *ΔThnag : : hph* mutant.

**Figure 1.**
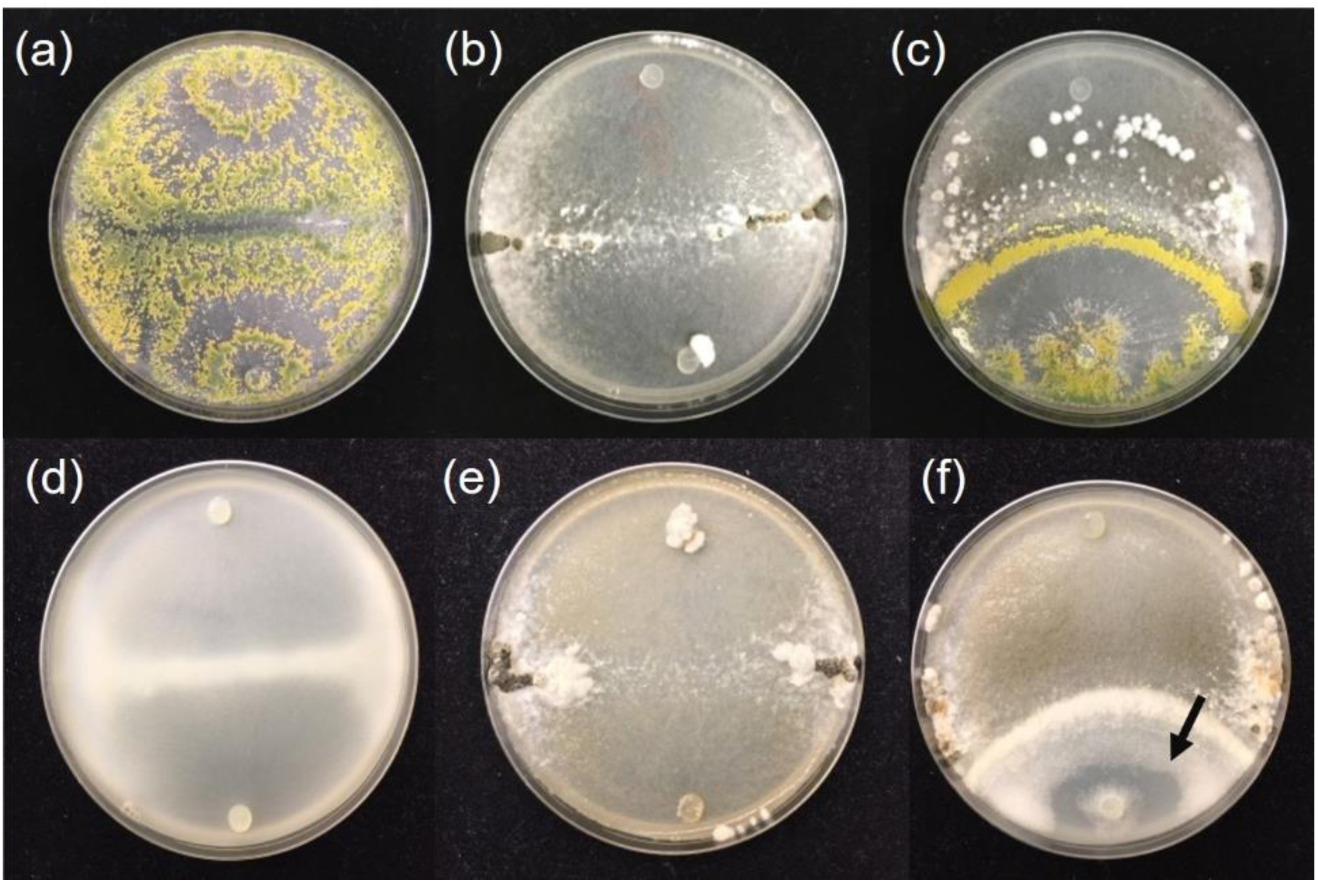
Dual-culture confrontation assays of (a) self-challenged *Trichoderma hamatum* GD12 strain; (b) self-challenged *Sclerotinia sclerotiorum* (c) co-culture of *S. sclerotiorum* (top) and *T. hamatum* GD12 (bottom); (d) self-challenged *T. hamatum* N-acetyl-β-glucosaminidase (*ΔThnag : : hph*) mutant; (e) self-challenged *S. sclerotiorum*; (f) co-culture of *S sclerotiorum* (top) and *T. hamatum ΔThnag : : hph* mutant (bottom).

Significant quantitative and qualitative changes in VOC production compared to self-challenged GD12 treatments were observed when *T. hamatum* GD12 was co-cultured with *S. sclerotiorum* (Figure 2; Table 1). Production of 6-pentyl-2H-pyran-2-one (6-PAP) dominated the headspace of GD12 co-cultured with *S. sclerotiorum*, with mean production of 6-PAP significantly greater in these co-cultures compared to self-challenged controls (p <0.001) (Table 1). Of the 36 compounds detected, eight of which were confirmed by co-injection, 22 were unique to GD12-*S. sclerotiorum* co-cultures, suggesting that the VOCs were either biosynthesised *de novo* in the presence of *S. sclerotiorum* or produced below detectable limits of the GC in GD12 monocultures, including 2-pentylfuran (Figure 2; Table 1).

**Figure 2.**
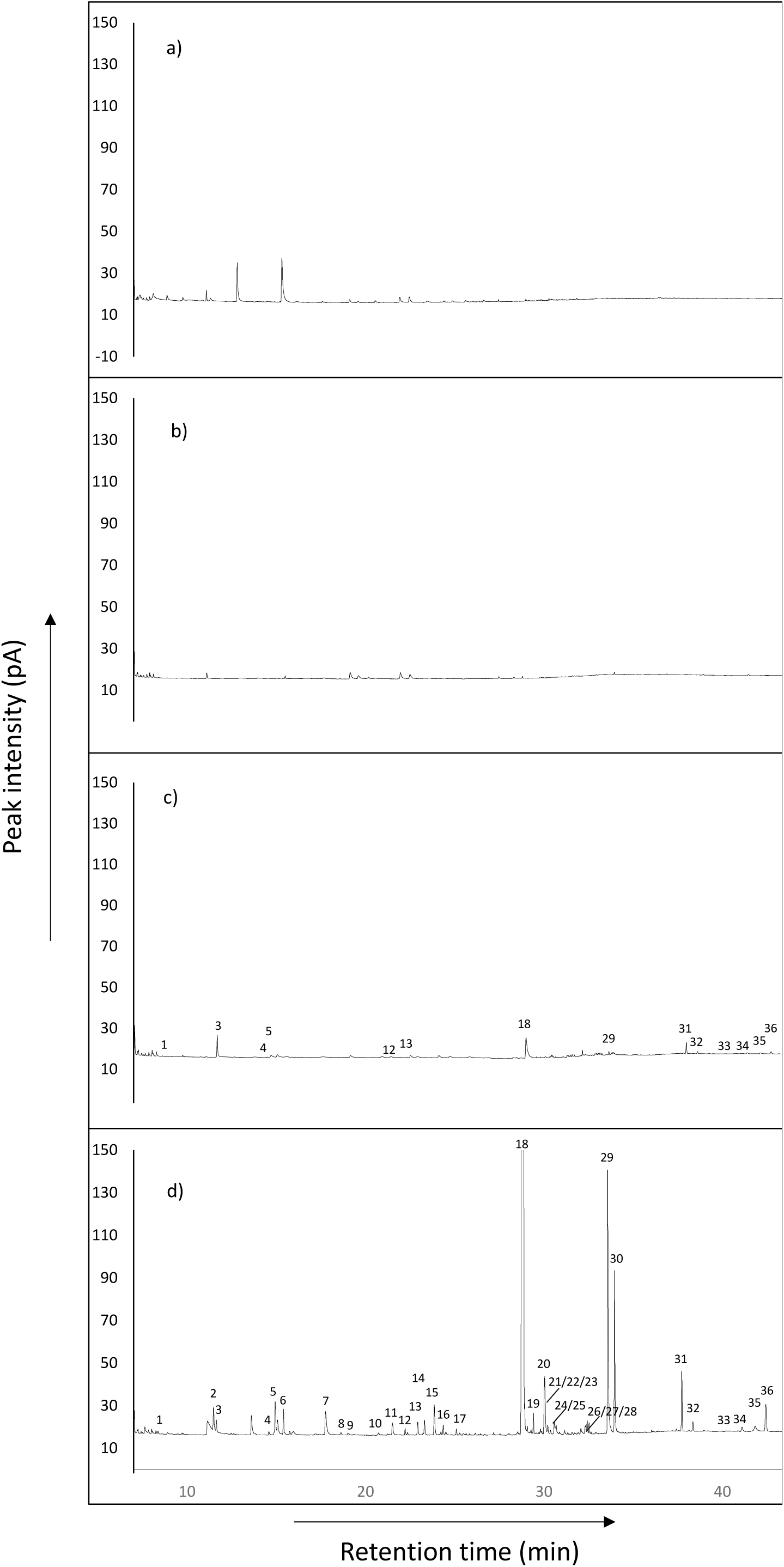
Representative gas chromatographic analysis of volatile organic compounds (VOCs) collected by air entrainment from 7-day old cultures of (a) uninoculated growth media (control), (b) self-challenged *S. sclerotiorum*, (c) self-challenged *T. hamatum* GD12, and (d) *T. hamatum* GD12 co-inoculated with *S. sclerotiorum*. For an explanation of peak numbers, see Table 1.

**Table 1.**
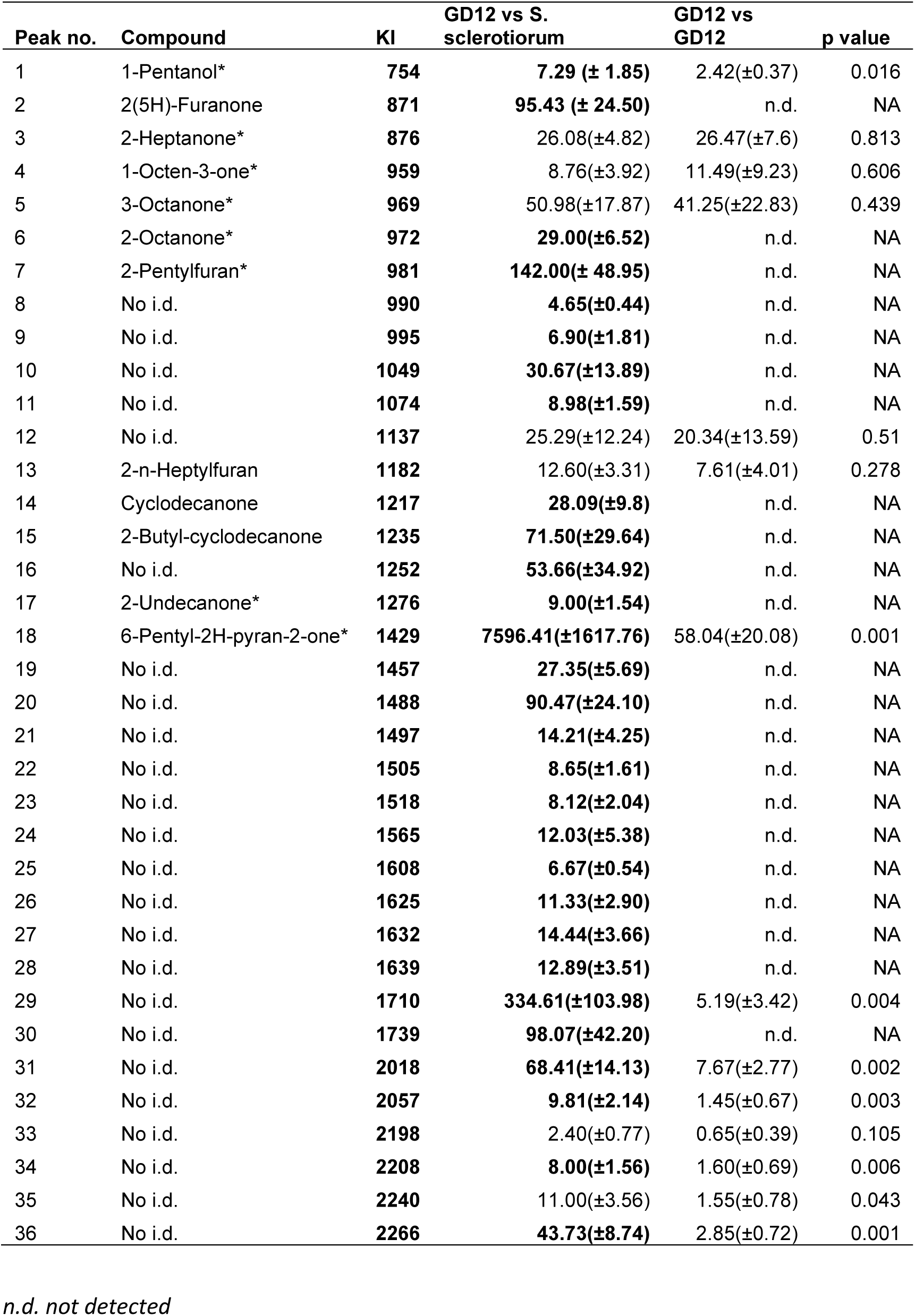
Composition of VOCs from dynamic headspace collections of 7-day old cultures of self-challenged *T. hamatum* GD12, or GD12 co-inoculated with *S. sclerotiorum* (n=4) (mean peak area ± SE). Data were analysed by Student’s t-test (p < 0.05). KIs on a non-polar HP-1 GC column.

As co-culturing *T. hamatum* GD12 with *S. sclerotiorum* led to significant increases in VOC production, the temporal dynamics of *T. hamatum* GD12 VOC production was investigated. For three compounds detected by SPME, peak induction was greatest at day 17 post-co-culture, each subsequently decreasing by day 24 (Figure 3). Each compound showed similar trends in production across the different timepoints. In GD12 co-cultured with *S. sclerotiorum* treatments, compound 1 (KI = 1252) was not detected in the headspace until day 6. By day 10 there was an increase in production, which further increased at day 17, subsequently decreasing to day 24. 6-PAP shows a similar trend. Compound 2 (KI = 1994), detected by day 10, increasing until day 17, and then decreasing by day 24.

**Figure 3.**
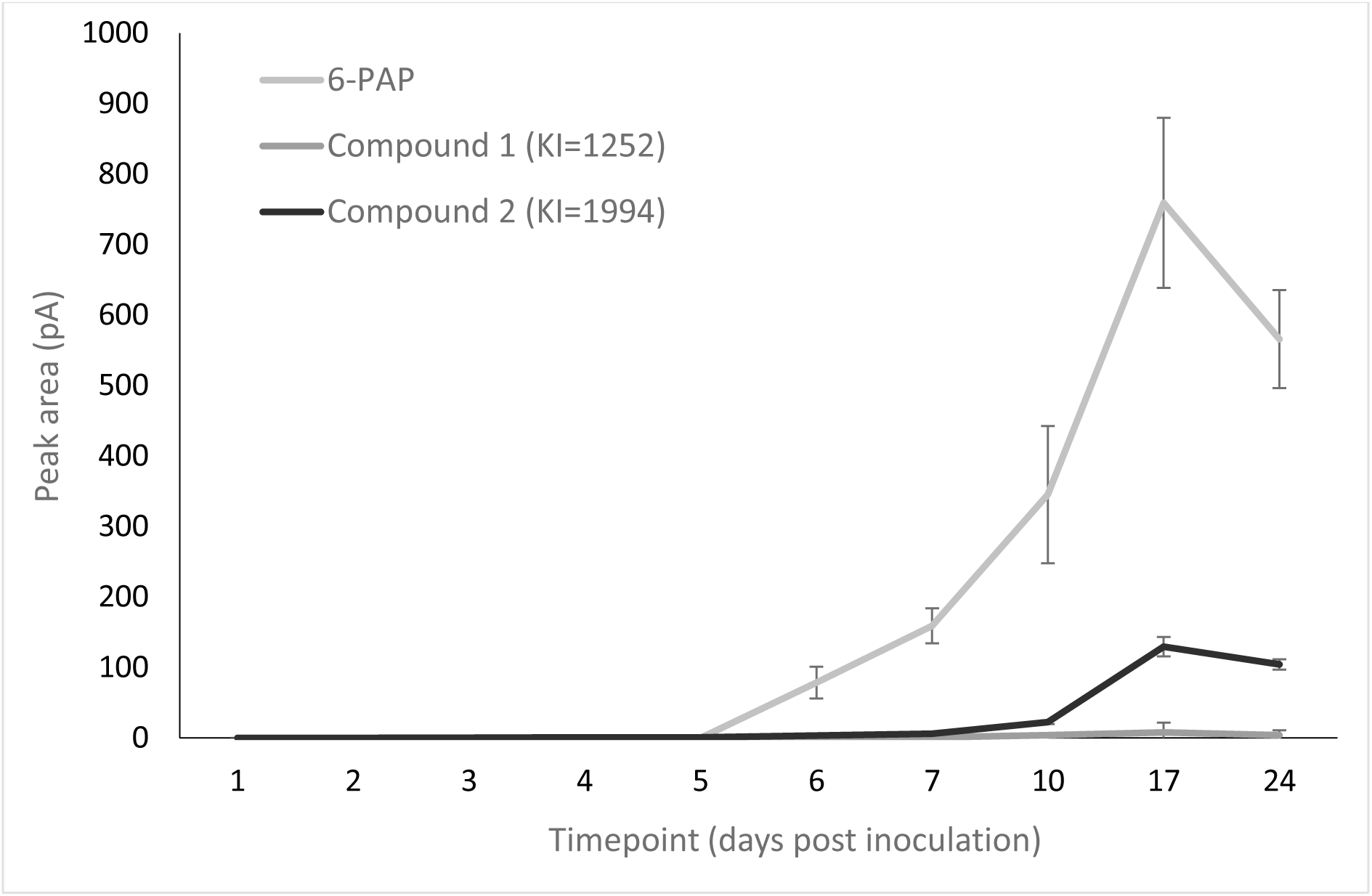
Production of VOCs in treatments of *Trichoderma hamatum GD12* and *S. sclerotiorurm* co-culture treatments over the course of 24 days for three VOCs. Bars represent the peak area value of each VOC (± SD) (n=3).

When co-cultured with *S. sclerotiorum*, quantitative and qualitative changes in VOC production relative to self-challenged *ΔThnag : : hph* treatments were observed, although unlike with GD12, most VOCs were either down-regulated during co-cultivation, or not significantly different to self-challenged controls (Figure 4). This provides a negative control to directly link unique GD12 VOC production to biocontrol. Four of the 10 VOCs detected are up-regulated in self-challenged *ΔThnag : : hph* treatments relative to *ΔThnag : : hph* co-cultured with *S. sclerotiorum*, and five were not significantly different to controls (Table 2). Notably, 6-PAP, showing the greatest induction in GD12 co-cultured with *S. sclerotiorum* treatments, was present in *ΔThnag : : hph* monocultures, but not in three out of four replicates in *ΔThnag : : hph* co-cultured *S. sclerotiorum* treatments. Four VOCs (KI = 1552; KI = 2016; KI =2055; KI = 2264) were significantly greater in self-challenged *ΔThnag : : hph* relative to co-culture treatments, and five were not significantly different. Taken together, these findings indicate a direct or indirect role for the *N*-acetyl-β-glucosaminidase enzyme in the induction of VOCs by *T. hamatum* GD12 during confrontation with *S. sclerotiorum*.

**Figure 4.**
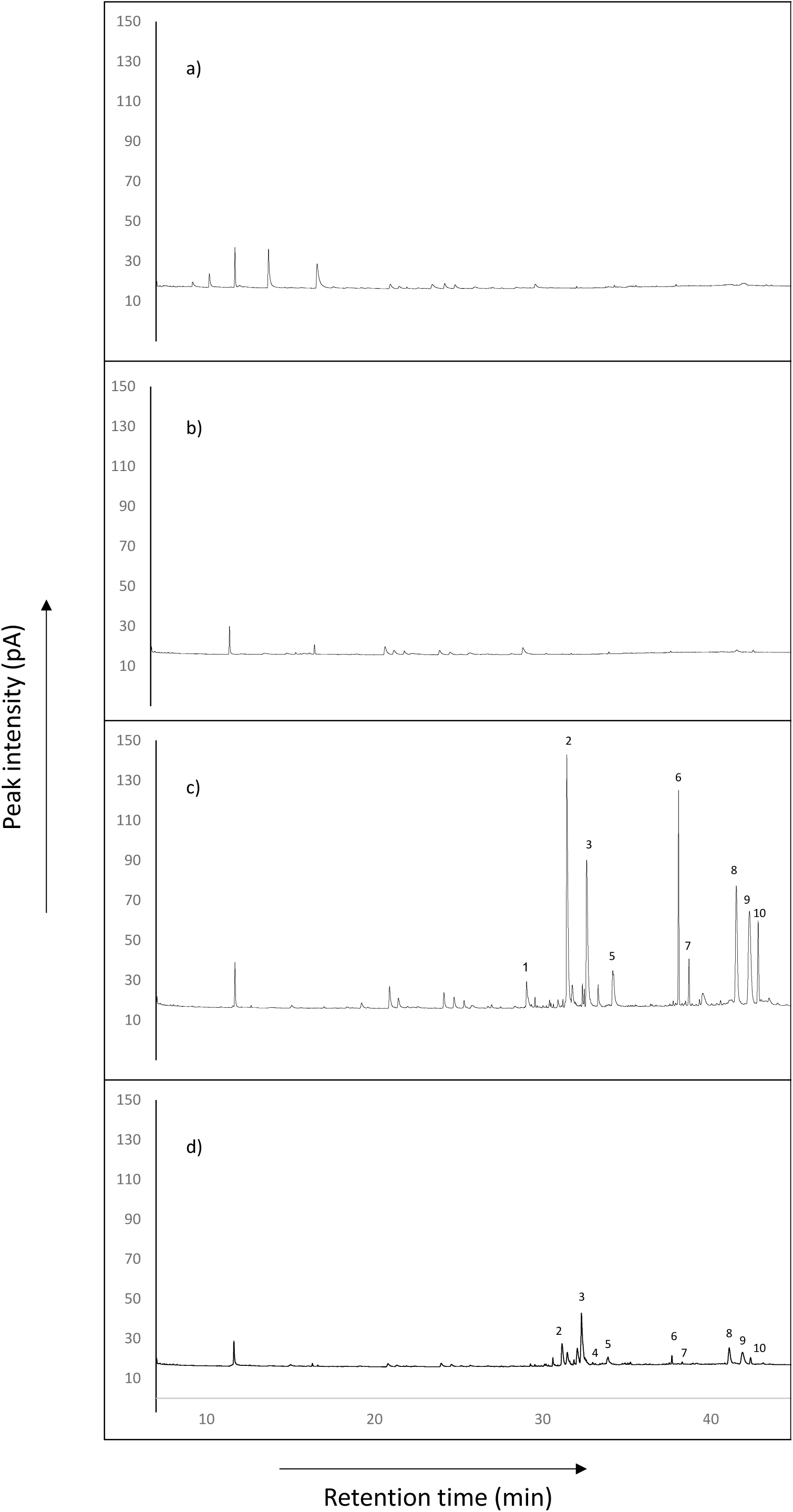
Representative gas chromatographic (on an HP-1 column) analysis of VOCs from dynamic headspace collections of 7-day old cultures of (a) uninoculated growth media (control), (b) self-challenged *S. sclerotiorum*, (c) self-challenged *T. hamatum ΔThnag : : hph*, and (d) *T. hamatum ΔThnag : : hph* co-inoculated with *S. sclerotiorum*.

**Table 2.** Composition of VOCs collected from dynamic headspace collections of 7-day old cultures of self-challenged *T. hamatum ΔThnag : : hph*, or *ΔThnag : : hph* co-inoculated with *S. sclerotiorum* (n=4) (mean peak area ± SE). Data were analysed by Student’s t-test (p < 0.05). KIs on a non-polar HP-1 GC column.

| Peak no. | KI | Compound | Mean (± SEM) | | |
| --- | --- | --- | --- | --- | --- |
|  |  |  | nag vs *S. sclerotiorum* | nag vs nag | p value |
| 1 | 1425 | 6-Pentyl-2H-pyran-2-one* | n.d. | 52.06 (±17.40) | NA |
| 2 | 1552 | No i.d. | 61.89(±21.1) | **793(±213.9)** | 0.049 |
| 3 | 1624 | No i.d. | 189.32(±69.91) | 526.03(±145.52) | 0.278 |
| 4 | 1692 | No i.d. | **1.9(±0.78)** | 0 |  |
| 5 | 1735 | No i.d. | 26.14(±9.06) | 161.8(±53.61) | 0.118 |
| 6 | 2016 | No i.d. | 51.09(±2.80) | **322.7(±66.90)** | 0.001 |
| 7 | 2055 | No i.d. | 3.37(±1.20) | **80.3(±16.74)** | 0.003 |
| 8 | 2208 | No i.d. | 74.87(±28.26) | 336.83(±78.090) | 0.15 |
| 9 | 2243 | No i.d. | 76.49(±28.95) | 367(±90.19) | 0.145 |
| 10 | 2264 | No i.d. | 22.31(±2.91) | **271.00(±67.83)** | 0.005 |

In preliminary experiments with *T. hamatum* VOCs and *S. sclerotiorum*, 6-PAP had no detectable inhibitory activity against *S. sclerotiorum* (F2,6=2.17, p = 0.195) (Figure S2) and was therefore excluded from further bioassays. The mycelial area of *S. sclerotiorum* differed significantly depending on the specific VOC applied at the highest dose (45.5 µM), indicative of differences in their inhibitory activities (F7,16= 171.36; p < 0.001; n= 3) (Figure 5). The mycelial area of *S. sclerotiorum* exposed to 1-pentanol was not significantly different to solvent control treatments (P > 0.05), and 2-heptanone treatments were not significantly different to 1-pentanol treatments (P > 0.05), so these compounds were not tested at reduced doses. 2-octanone and 2-undecanone demonstrated similar levels of inhibition, while 2-octanone was significantly more inhibitory than 3-octanone. Strikingly, 1-octen-3-one demonstrated effectively complete growth of inhibition of *S. sclerotiorum*. Importantly, this antifungal activity was effective against other economically fungal pathogens (*Botrytis cinerea*, *Pyrenopeziza brassicae* and *Gaeumannomyces tritici*) (Figure S3).

**Figure 5.**
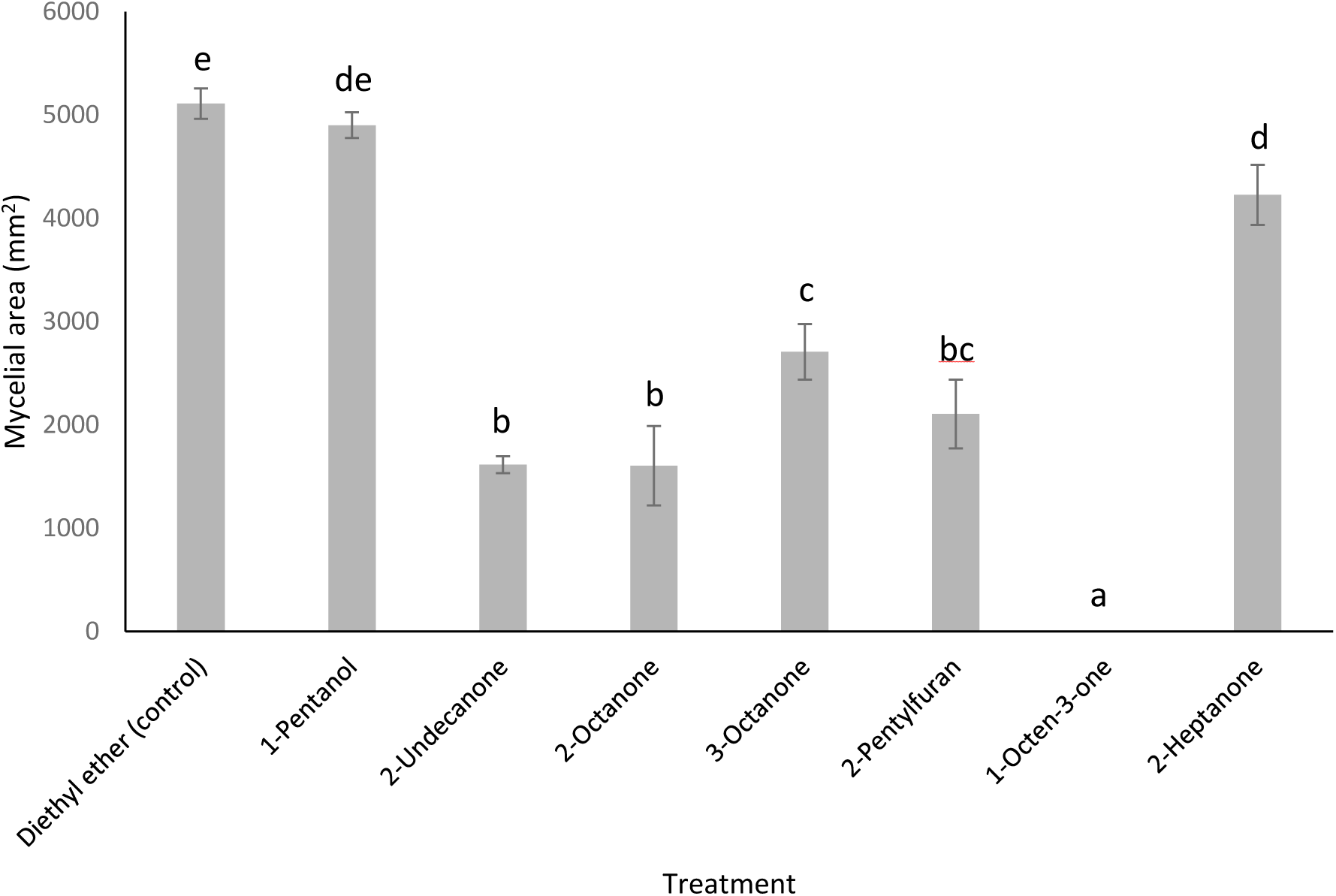
Antifungal activities of selected VOCs on the growth of *S. sclerotiorum*. *S. sclerotiorum* was incubated with selected VOCs at 45.5 µM doses and the inhibition rates were calculated relative to control plates (exposed to diethyl ether alone) after 3 days. Bars represent the mean mycelial area of *S. sclerotiorum* upon exposure to each VOC (± SD) (n=3). Different letters indicate significant differences between treatments according to Tukey’s multiple comparisons test (p < 0.05). Y axis represents mycelial area (mm^2^).

Of the tested VOCs, the five demonstrating the most significant inhibition were selected for further study at reduced doses, and all demonstrated significant inhibition when applied at reduced doses; 1-octen-3-one: (F7,16= 246.03; p < 0.001); 2-octanone: (F3,8= 42.29; p < 0.001); 3-octanone: (F4,10= 31.74; p < 0.001); 2-pentylfuran: (F5,12= 119.2; p < 0.001) and 2-undecanone: (F7,16= 91.65; p < 0.001) (Figure 5). 2-octanone had a minimum inhibitory dose of 11.125 µM, 3-octanone, 2-pentylfuran had a minimum inhibitory dose of 4.55 µM and 2-undecanone, 2.275 µM (Figure 6). The compound showing the greatest inhibition, 1-octen-3-one, was still significantly inhibitory at a 100-fold dilution (0.445 µM).

**Figure 6.**
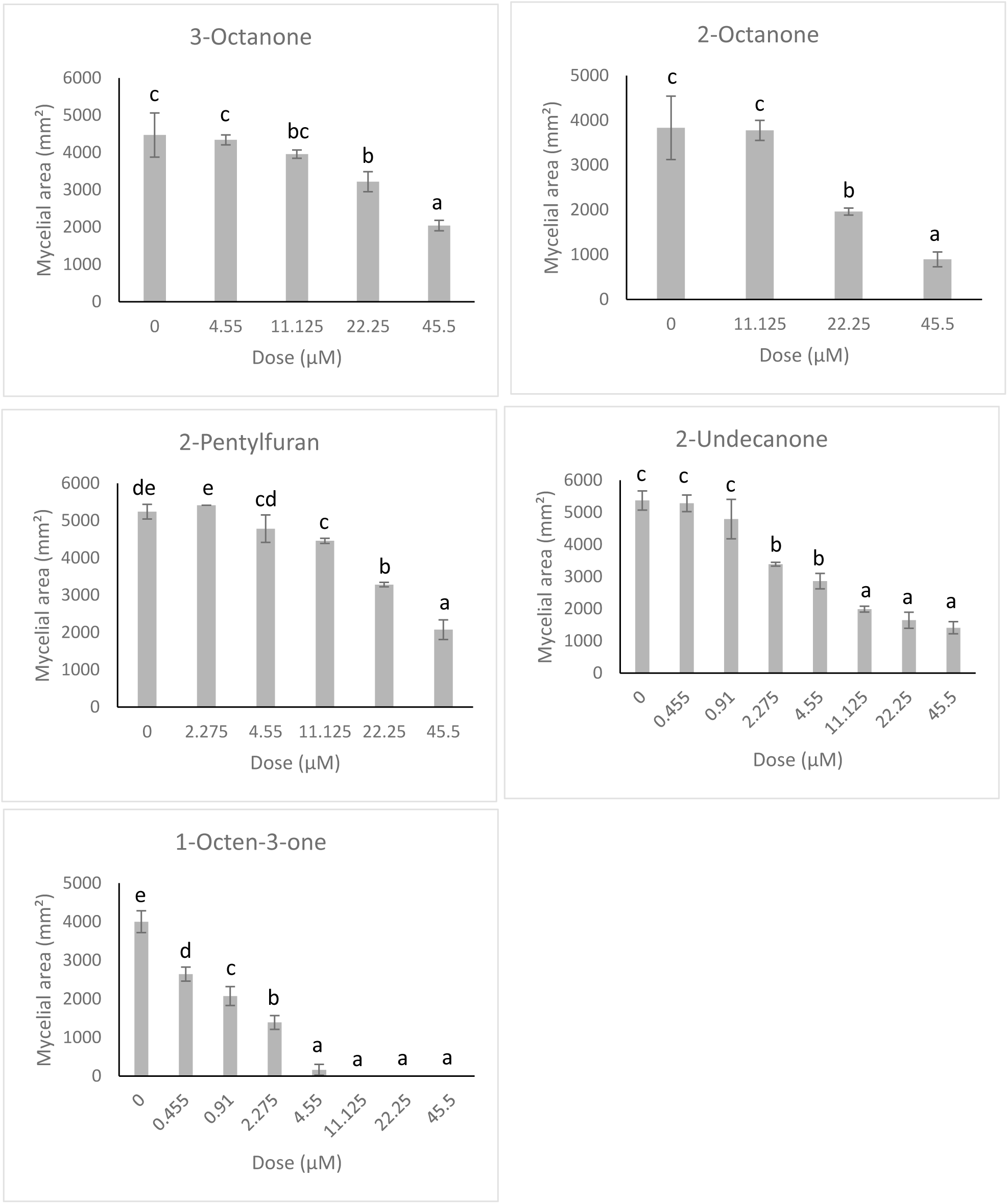
Antifungal activities of selected VOCs on the growth of *S. sclerotiorum*, at reduced doses. Bars represent the mean mycelial area of *S. sclerotiorum* upon exposure to each VOC (± SD) (n=3). Different letters indicate significant differences between treatments according to Tukey’s multiple comparisons test (p < 0.05). Y axis represents mycelial area (mm^2^).

To establish potential structural moieties required for antifungal activity of 1-octen-3-one against *S. sclerotiorum*, compounds with similar structural features to both 1-octene, 3-octanone and (*RS*)-1-octen-3-ol were tested (Figure S4), revealing significant differences in inhibitory activities (F4,10= 114.44; p < 0.001) (Figure 7). 1-octene demonstrated no significant inhibition of *S. sclerotiorum* relative to solvent controls (p > 0.05), whereas both 3-octanone and (*RS*)-1-octen-3-ol showed significant inhibition of *S. sclerotiorum* relative to controls (p < 0.05). However, only 1-octen-3-one demonstrated 100% inhibition. When *S. sclerotiorum* was removed from the shared atmosphere with 1-octen-3-one, fungal growth of the pathogen was not restored 4 weeks after removing the pathogen from the headspace (Figure 8). This is consistent with 1-octen-3-one having fungicidal activity at the tested dose.

**Figure 7.**
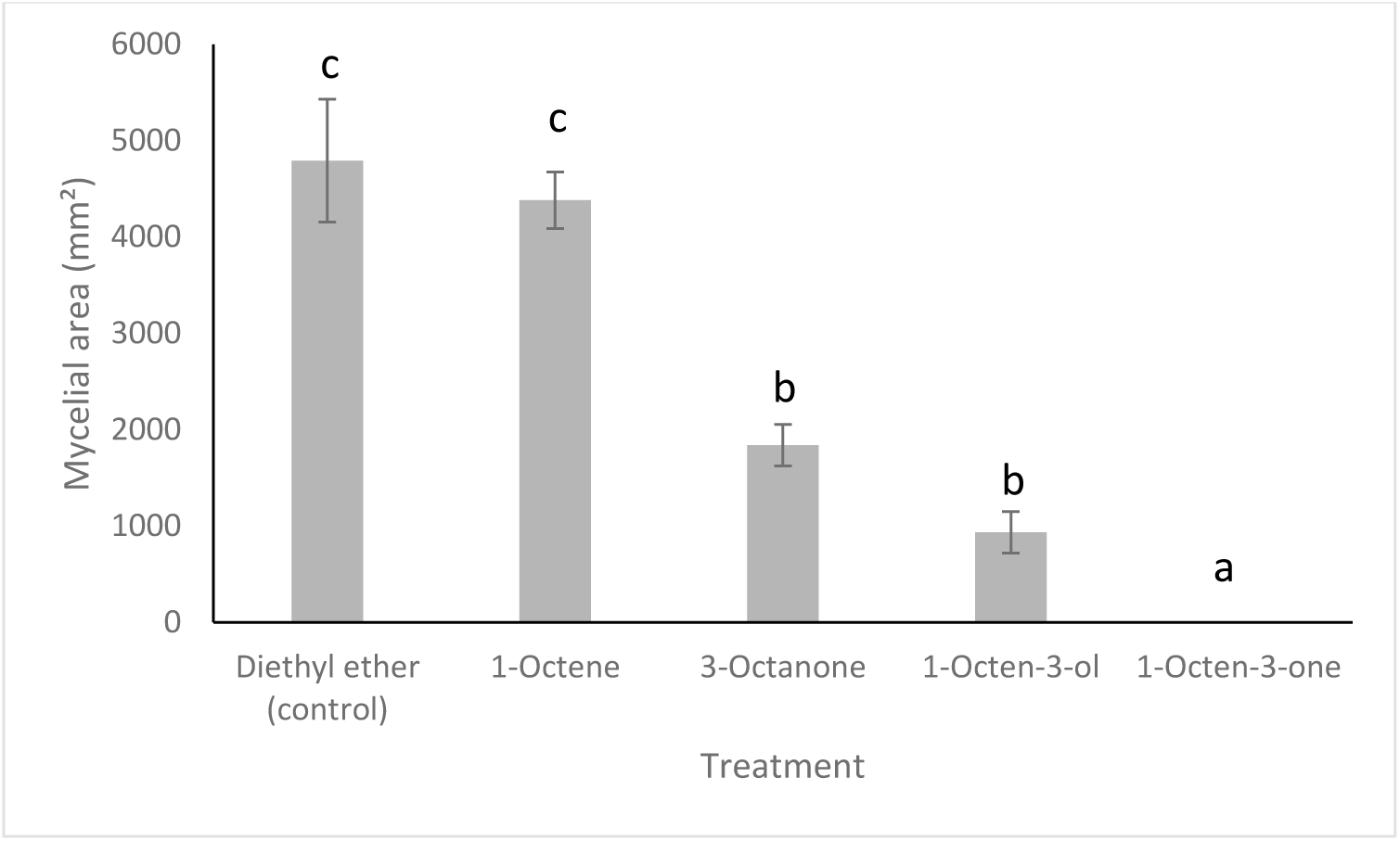
Antifungal activities of selected VOCs on the growth of *S. sclerotiorum*, representing individual structural components of 1-octen-3-one, at 45.5 µM. Bars represent the mycelial area of *S. sclerotiorum* upon exposure to each VOC (± SD) (n=3). Different letters indicate significant differences between treatments according to Tukey’s multiple comparisons test (p < 0.05).

**Figure 8.**
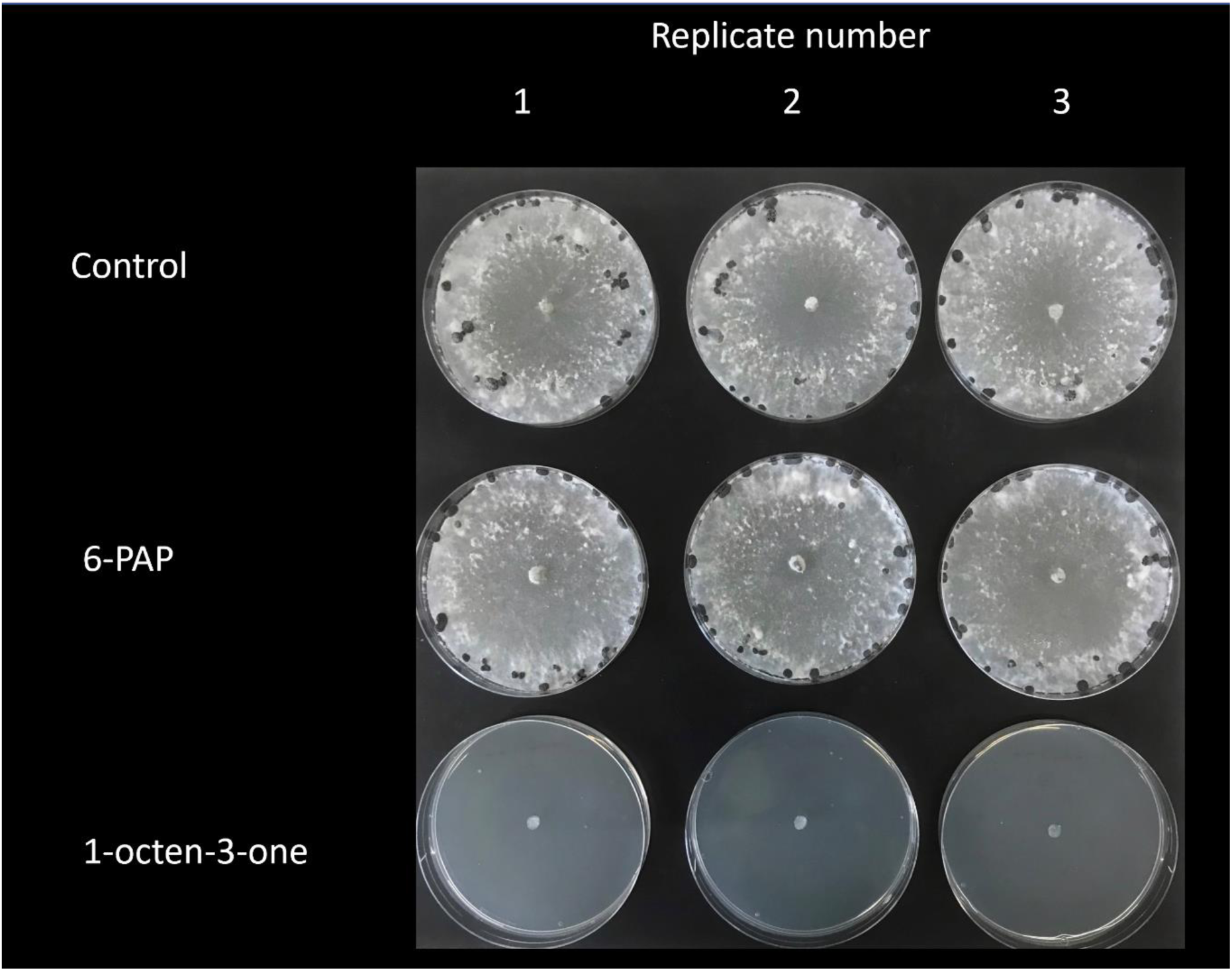
Antifungal activities of 6-PAP and 1-octen-3-one, on *Sclerotinia sclerotiorum* at 45.5 µM.

## Discussion

In this study, we showed induction of VOC production by *T. hamatum* GD12 during co-culture with the fungal pathogen *S. sclerotiorum*. This included VOCs not produced by *T. hamatum* when grown axenically. This VOC induction was not observed during the interaction between *S. sclerotiorum* and the *ΔThnag : : hph* mutant, suggesting a role of the *N*-acteyl-β-glucosaminidase enzyme in the direct or indirect facilitation of VOC induction. Whilst several VOCs were also produced in self-challenged GD12 controls, the stimulation of VOC production in co-cultures indicates a *de novo* biosynthesis in the presence of the pathogen, some of which we demonstrate possess an antifungal role against *S. sclerotiorum*.

Many studies investigating VOC production from *Trichoderma* species utilise axenic fungal growth, and VOCs are predominantly assigned to alcohols, ketones, alkanes, furans, mono-and-sesquiterpenes (Jeleń *et al*., 2014a). Several low molecular weight compounds reported here as being produced by *T. hamatum* GD12 have been previously identified from other *Trichoderma* species, including 3-octanone (Nemčovič *et al*., 2008; Stoppacher *et al*., 2010; Jeleń *et al*., 2014a; Estrada-Rivera *et al*., 2019; Speckbacher *et al*., 2020; Silva *et al*., 2021) and 2-octanone (Jeleń *et al*., 2014b; Estrada-Rivera *et al*., 2019; Speckbacher *et al*., 2020). To our knowledge, this is the first report of 1-octen-3-one being produced by *T. hamatum*, although it was identified from *T. virens* (Li *et al*., 2018), and a range of other fungi (Pennerman *et al*., 2022). 6-PAP is a characteristic *Trichoderma* VOC (Mendoza-Mendoza *et al*., 2024), which produces a coconut aroma (Reithner *et al*., 2007; Stoppacher *et al*., 2010; Jeleń *et al*., 2014a; Garnica-Vergara *et al*., 2016; Estrada-Rivera *et al*., 2019; Baazeem *et al*., 2021; Silva *et al*., 2021). When comparing the *T. hamatum* GD12 VOCs identified via co-culture with other studies, only 6-PAP has been previously reported (Jeleń *et al*., 2014a; Baazeem *et al*., 2021). However, directly comparing volatile diversity across other studies should be undertaken with caution as such studies employ different volatile sampling techniques, which can introduce biases for certain compounds. For example, while many studies deploy SPME for headspace sampling, the diversity of fungal volatiles recovered depends on the type of fibre used (Stoppacher et al., 2010; Jeleń et al., 2014). Growth conditions of cultures will also likely vary across studies, which can influence volatile production from *Trichoderma*, including age of cultures (Lee *et al*., 2015), relative humidity and temperature of growth conditions (Polizzi *et al*., 2011), as well as media composition (Zhang *et al*., 2014; González-Pérez *et al*., 2018). Intraspecific differences in volatile production have been observed for *T. hamatum*, as well as within other *Trichoderma* species. This likely relates to fungal evolution/adaptation to different geographical regions or ecological niches from which they were isolated (Jeleń et al., 2014; Lee et al., 2016), but also highlights the power of geographical/niche adaptation to drive the evolution of novel antifungals . Taken together, a range of factors can account for variation in VOC production by *Trichoderma*, both inter- and intra-specifically.

Co-cultivation with *S. sclerotiorum* revealed significant quantitative and qualitative changes in volatile production compared to the VOCs produced by *T. hamatum* GD12 in self-challenged controls. This induction was greatest 17 dpi, consistent with other studies (Hynes et al., 2007). 2-Pentylfuran, which was biosynthesised *de novo* in response to the interaction with *S. sclerotiorum*, has previously been reported from *T. hamatum* during axenic culture (Jeleń *et al*., 2014a). Many of the upregulated compounds in our study are of the sesquiterpene-like class, which have previously been observed during physical fungal-fungal interactions (Hynes *et al*., 2007a; Guo *et al*., 2019a; Rajani *et al*., 2021), including *T. hamatum* when challenged with the ectomycorrhizal fungus *Laccaria bicolor* (Guo *et al*., 2019b) and *Trichoderma* in confrontation with *Sclerotium rolfsii* and *Macrophomina phaseolina* (Sridharan *et al*., 2020). Sesquiterpenes are a well-known class of compounds involved in chemical signalling, and have been isolated across a range of *Trichoderma* species, including *T. hamatum* (Ma *et al*., 2021), *T. brevicompactum* (Shi *et al*., 2020), *T. virens* (Shi *et al*., 2018, 2021; Hu *et al*., 2019), *T. longibrachiatum* (Du *et al*., 2020; Wang *et al*., 2022), *T. asperellum* (Ding *et al*., 2012), and *T. citrinoviride* (Liu *et al*., 2020). Many of these sesquiterpene-like compounds possess antifungal activities against a range of phytopathogenic fungi, bacteria and marine phytoplankton. As well as their antimicrobial roles, microbial sesquiterpenes have a range of other biological activities including signalling, host growth promotion and defence (Avalos *et al*., 2022). The upregulation of unknown sesquiterpenes during co-culture of *T. hamatum* and *S. sclerotiorum* could indicat a biological role for these compounds, and future work aims to isolate and identify these compounds to determine their role in the antagonistic response against *S. sclerotiorum*, and potential for integrating into biocontrol strategies.

6-PAP dominated the VOC profile of *T. hamatum* GD12 co-cultured with *S. sclerotiorum*, relative to self-challenged *T. hamatum* GD12 cultures, corroborating previous work which found significant increases in 6-PAP production when *T. harzianum* was co-inoculated with *R. solani* (Serrano-Carreón *et al*., 2004; Flores *et al*., 2019). Whilst significant increases in 6-PAP production by *T. hamatum* GD12 in co-culture were observed, no antifungal activity was observed when 6-PAP was applied in the inverted plate assay setup (Figure S2). However, several studies demonstrate an inhibitory role for 6-PAP against a range of fungal pathogens, including *Fusarium* species (Scarselletti and Faull, 1994; El-Hasan *et al*., 2008; Jeleń *et al*., 2014a; Rao *et al*., 2022), *Botrytis cinerea* (Pezet *et al*., 1999), *Cylindrocarpon destructans* (Jin *et al*., 2020), and *Rhizoctonia solani* (Scarselletti and Faull, 1994), when the compound was in contact with the pathogens. These studies indicate that 6-PAP may require direct contact for effective antifungal activity. As well as 6-PAP upregulation, 2-octanone production was significantly upregulated in co-culture treatments, indicative of *T. hamatum* - *S. sclerotiorum* antagonism. 2-octanone upregulation has also been observed during the interaction between the fungal pathogen *Setophoma terrestris* and the beneficial soil bacteria *Bacillus subtilis* (Albarracín Orio *et al*., 2020), as well as the interaction between *T. atroviride* and *F. oxysporum* (Speckbacher *et al*., 2021), indicating a broader spectrum role for this compound in antagonistic fungal interactions.

Many of the VOCs produced by *T. hamatum* GD12 during self-challenge and in co-cultures with *S. sclerotiorum* show significant antifungal activity against *S. sclerotiorum*. The antifungal role of 2-octanone demonstrated here is in agreement with the inhibition of the soil fungal pathogen *Setaphoma terrestris*, which could indicate broad-spectrum inhibitory activity against a range of fungal pathogens (Albarracín Orio *et al*., 2020). Similarly, 2-heptanone has previously shown significant inhibition against *Curvularia lunata* (Xie *et al*., 2020) and *Alternaria solani* (Zhang *et al*., 2020). Here, we report only moderate antifungal activity of 2-heptanone at the highest tested dose relative to other compounds, which may relate to differences in doses tested across the studies, or specificity in the antifungal activity of 2-heptanone against different pathogenic species. Alternatively, it may reflect that anti-fungal role could be derived from 2-octanone via further modifications that many occur in a more complex soil microbiome, as opposed to our two-component experimental system. Specificity of antifungal activity has been observed for 2-undecanone, which has shown an inhibitory role against *Verticillium dahlia*, *F. oxysporum*, *B. cinerea* and *Monilinia* spp., but not *Penicillium* spp. (Calvo *et al*., 2020) or *Rhizopus stolonifer* (Carter-House *et al*., 2020). Having identified a range of antifungal compounds, determining the modes of action of the antifungal activities against *S. sclerotiorum* is an important next step. Furthermore, whilst the inhibitory properties of several *T. hamatum* VOCs have been demonstrated against *S. sclerotiorum*, it is important to establish that compounds at their inhibitory doses do not have phytotoxic effects. Interestingly, 2-pentylfuran, which was upregulated in the presence of *S. sclerotiorum* and showed an antifungal role against the pathogen, has also demonstrable plant growth-promoting capabilities (Zou *et al*., 2010). Thus, 2-pentylfuran could potentially be a promising candidate to replace synthetic chemical inputs due to its ability to inhibit fungal pathogens without compromising plant growth.

To our knowledge, this is the first report demonstrating an antifungal role for 1-octen-3-one against a pathogen. When structurally related compounds (1-octene, 3-octanone, (*R,S*)-1-octen-3-ol) were tested for their antifungal activity at equivalent doses, 3-octanone and (*RS*)-1-octen-3-ol demonstrated significant inhibition of *S. sclerotiorum*, whereas absolutely no growth of *S. sclerotiorum* occurred when exposed to 1-octen-3-one. It is thus possible that fungicidal activity may be enhanced via Michael-type acceptance by the α,b-unsaturated carbonyl structure within the latter compound (Li *et al*., 2019). Several of the strobilurin class of fungicides (fungicides derived from *Strobilurus spp.*, Nofiani *et al*., 2018) also contain a conjugated ketone and alkene moiety. Findings here are contrary to those reported by Xiong *et al*. (2017), who found significant inhibition of *F. tricinctum* and *F. oxysporum* treated with 1-octen-3-ol, but no inhibition when fungi were treated with 1-octen-3-one (Xiong *et al*., 2017). However, VOCs were administered differently in each experiment, making cross-comparison difficult. VOCs tested by Xiong et al. were supplemented into growth medium and in direct contact with *Fusarium* species, whereas tested VOCs here were physically separated from *S. sclerotiorum,* suggesting that the antifungal effect of 1-octen-3-one works at a distance. An important consideration is that, in both studies, (*RS*)-1-octen-3-ol was tested as a racemic mixture, although previous work has shown chirality can impact its antifungal activity (Yin *et al*., 2019). Whilst 1-octen-3-one shows an inhibitory role against *S. sclerotiorum*, future work should determine the role of the compound on plant growth. For example, 1-octen-3-one exposure significantly inhibits *Arabidopsis* growth and development (Lee *et al*., 2019), therefore future work should focus on determining which dose of 1-octen-3-one can inhibit *S. sclerotiorum* without compromising plant growth. Moreover, as seen with antimicrobial drugs, a mixture of antifungals may have synergistic activities, yet overall reduced potential phytotoxic effects, enabling the incorporation of 1-octen-3-one into biocontrol strategies. As compounds identified here have *in vitro* antagonism against *S. sclerotiorum*, immediate priorities will be to test these compounds against *S. sclerotiorum*, individually and in combinations, using peat microcosms under glasshouse conditions. These data will inform future open field trials, to examine their biological activities under more agriculturally relevant conditions.

*Trichoderma* spp. recognise plant pathogenic fungi when their lytic enzymes, including *N-*acetyl*-β-*D*-* glucosaminidase, release small diffusible components from fungal cell walls (Druzhinina *et al*., 2011). These components can then bind G-Protein Coupled Receptors (GPCRs) on the cell surface of *Trichoderma*, activating a downstream signalling cascade leading to the expression of secondary metabolite biosynthesis genes, potentially including genes linked to volatile production. As *ΔThnag : : hph* cannot produce the chitinase enzymes required to break down fungal cell walls, theoretically no breakdown products from *S. sclerotiorum* would bind to the GPCRs of *T. hamatum*, preventing elicitation of the downstream signalling cascade and activation of secondary metabolite biosynthesis genes. Significant reduction in 6-PAP production by the *ΔThnag : : hph* mutant during co-culture with *S. sclerotiorum* may also suggest a role for *N-*acetyl*-β*-D-glucosaminidase in 6-PAP production, which several studies have previously demonstrated. For example, deletion of the *tga1* gene, encoding the α-subunit of a heterochromatic G-protein 1 from *T. atroviride*, led to both a reduction in *N-*acetyl*-β*-D-glucosaminidase activity in mutant strains and an 8-fold reduction in 6-PAP production, relative to controls (Reithner *et al*., 2005). Similarly, deletion of the gene encoding mitogen-activated protein kinase (*tmk1*) led to a 1.6-fold increase in production of 6-PAP and the enhancement of *nag1* expression relative to the parent strain (Reithner *et al*., 2007). The reduction in 6-PAP production by *Δthnag : : hph* may also explain why the mutant loses its antagonistic activity against *S. sclerotiorum* in peat microcosms (Ryder *et al*., 2012).

In conclusion, this study suggests a role for volatile chemical signalling during the antagonistic response of *T. hamatum* GD12 against *S. sclerotiorum* and shows that certain *Trichoderma*-derived VOCs play an inhibitory role against the pathogen. Specifically, we identify1-octen-3-one as a potential novel antifungal VOC. Further glasshouse and field tests with antifungal compounds identified here are required to determine whether they inhibit pathogens at a larger scale under more agriculturally relevant conditions. Whilst several of these compounds have been identified here, or previously described, many *T. hamatum* compounds upregulated on confrontation with fungal plant pathogens remain to be identified and characterised.

## Author contributions

Conceptualisation: MB, MG, CT. Data curation; GT, JV, JC, MB. Formal analysis; GT, JC, JV, MB, DW. Funding acquisition; MB, CT, MG. Investigation; MB, GT, MG, CT, JV, DW. Methodology; MB, GT, MG, CT, JS. Project administration; MB, MG, CT. Original draft; GT, MB, DW. Reviewing and editing; GT, MB, DW, JV, CT, MG, JS, JC.

## Acknowledgements

Rothamsted Research receives strategic funding from the Biotechnology and Biological Sciences Research Council of the United Kingdom (BBSRC). We acknowledge support from the Growing Health Institute Strategic Programme (BB/X010953/1; BBS/E/RH/230003A). The work formed part of the Rothamsted Smart Crop Protection (SCP) strategic programme (BBS/OS/CP/000001) funded through BBSRC’s Industrial Strategy Challenge Fund. GT’s PhD studentship was funded by a Biotechnology and Biological Sciences Research Council (BBSRC) Southwest Doctoral Training Programme award (project no. 1622285). The authors would also like to thank Jon West and Kevin King for providing *Sclerotinia sclerotiorum* and *Botrytis cinerea*, Lisa Humbert for providing *Pyrenopeziza brassicae*, and Javier Palma-Guerrero for providing *Gaeumannomyces tritici*.

