## Supplementary materials for "Inducible volatile chemical signalling drives antifungal activity of *Trichoderma hamatum* GD12 during confrontation with the pathogen *Sclerotinia sclerotiorum*"

Supplementary information

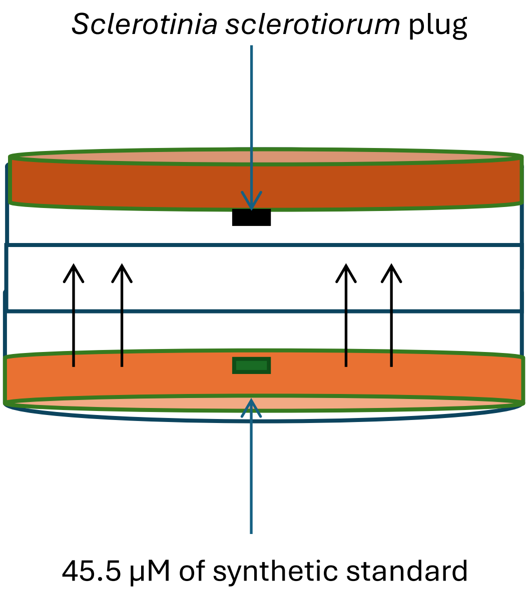


Figure S1 | Inverted plate bioassay set up

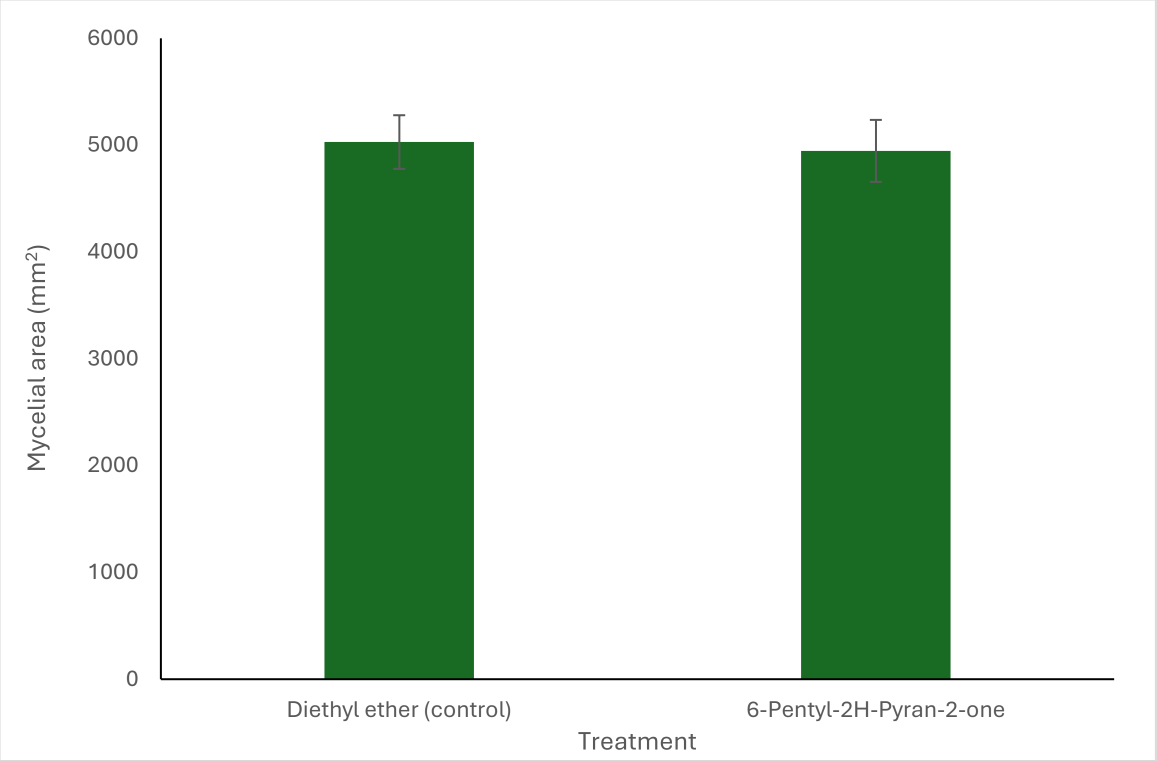


Figure S2 | Assessment of the antifungal activity of 6-Pentyl-2H-pyran-2-one (6-PAP) on the growth of *S. sclerotiorum*. *S. sclerotiorum* was incubated with 6-PAP at 45.5 µM doses, and the inhibition rates were calculated relative to control plates (exposed to diethyl ether alone) after 3 days. Bars represent the mean mycelial area of *S. sclerotiorum* upon exposure to each VOC (± SD) (n=3).

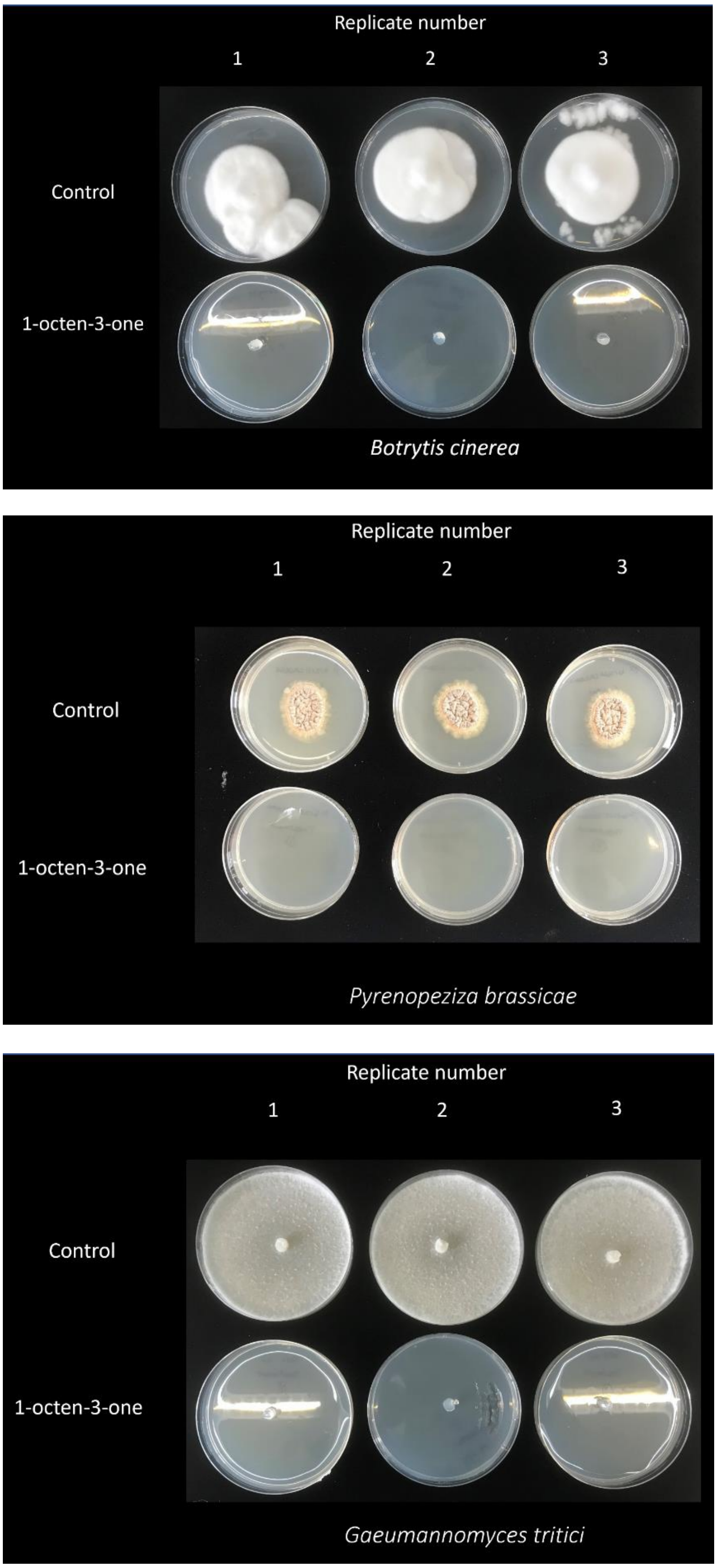


Figure S3 | Antifungal activities of 1-octen-3-one on the growth of *Botrytis cinerea*, *Pyrenopeziza brassicae* and *Gaeumannomyces tritici*. Fungal pathogens were incubated with selected VOCs at 45.5 μM doses for 3 days.

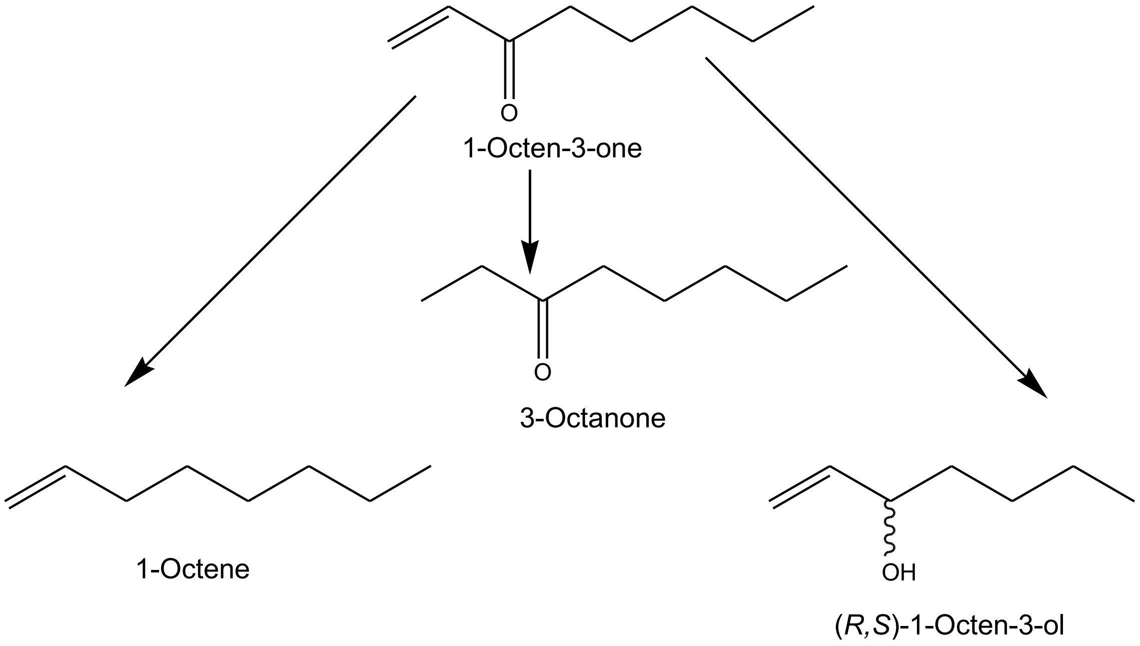


Figure S4 | Compounds which make up the individual structural components of 1-octen-3-one.
